# Preclinical safety evaluation of continuous UV-A lighting in an operative setting

**DOI:** 10.1101/2022.06.17.496643

**Authors:** Rachael Guenter, Rui Zheng-Pywell, Brendon Herring, Madisen Murphy, Kevin Benner, J. Bart Rose

**Affiliations:** Department of Surgery, University of Alabama at Birmingham, Birmingham, Alabama, USA; GE Current, a Daintree company, East Cleveland, Ohio, USA

## Abstract

**Background:** Germicidal ultraviolet (UV-C) light has been shown as an effective modality for disinfection in laboratory settings and in the operative room. Traditionally, short-wavelength UV-C devices, which have previously been shown to cause DNA damage, are utilized only for disinfection in pre- and post-operative settings and are not continuously active during operations. Continuous use of intraoperative UV light can potentially decrease pathogens and subsequent surgical site infections (SSIs), which arise in approximately 5-15% of operative cases. SSIs are a significant determinant of patient morbidity, readmission rates, and overall cost. Therefore, a method of UV light disinfection with a low risk of DNA damage is needed so that greater antimicrobial protection can be afforded to patients during the entirety of their surgical procedures. A new disinfection device that harnesses longer-wavelength UV-A light to disinfect the surgical field throughout the entirety of the procedure, including pre- and post-operation, has been developed.

**Methods:** This study aimed to determine if intraoperatively administered UV-A light was safe, as defined by the minimal presence of DNA damage and safe amounts of reflection upon medical personnel. Using *in vitro* models, we examined the differential impacts of UV-C and UV-A light on DNA damage and repair pathways. In a murine model, we looked at the difference in production of DNA damage photoproducts between UV-A and UV-C exposure.

**Results:** Our results show UV-A light does not induce a significant amount of DNA damage at the cellular or tissue level. Furthermore, a preclinical porcine study showed that surgical personnel were exposed to safe levels of UV-A irradiance from an overhead UV-A light used during an operation. The amount of UV-A transmitted through surgical personal protective equipment (PPE) also remained within safe levels.

**Conclusions:** In conclusion, we found that UV-A may be a safe for intraoperative use.

## INTRODUCTION

Approximately 30 million patients undergo surgical procedures each year in United States, and up to 5% of these patients will experience surgical site infections (SSIs), with infection rates as high as 15% for some operations (1). SSIs are significant determinants of morbidity, readmission, and cost to the healthcare system (2). Studies have also shown that patients who develop an SSI face greater financial burden from surgery when compared to uninfected patients (3). Additionally, patients with an SSI experienced prolonged hospitalization stays, which contributed to negative outcomes on their physical and mental health (3). The need for surgical procedures continues to increase, highlighting the urgent need for better SSI prevention practices to reduce medical complications and exacerbated costs (3,4). Several interventions have been proven to reduce SSIs, including appropriate antibiotic use, normothermia maintenance during procedures, and minimizing intra-abdominal bowel contamination (1,5). Despite these interventions, the rate of SSIs continues to rise globally (6).

The use of germicidal ultraviolet (UV) light has been shown to be an effective modality for disinfection of inanimate entities, such as room air and surfaces (7–9). Ultraviolet-C (UV-C) radiation, encompassing wavelengths ranging from 100 to 280 nm, has been used in healthcare settings to disinfect surfaces in operating rooms, hospital bed rooms, and ambulances (9–12). UV-C can also successfully sterilize the personal protective equipment (PPE) of healthcare workers (13). The germicidal properties of UV-C are attributed to the disruption of nucleic acids following exposure (13,14). Double bonds between the carbon atoms found in pyrimidines and purines are destabilized by UV-C, leading to the formation of dimers in RNA and DNA (15). Cyclobutyl pyrimidine dimers (CPD) and pyrimidine (6-4) pyrimidone photoproducts (6-4PP) are photoproducts that result from UV-C-induced DNA damage, and these photoproducts interrupt both the cell cycle and DNA replication (9,16,17). This cellular damage ultimately kills bacteria and causes viral inactivation (9,18–20).

Exposure of UV-C radiation can also cause harmful effects in humans; more specifically, negative effects on the skin and eyes have been reported after UV-C exposure (9). The ability of UV-C to induce DNA damage is carcinogenic (21). In fact, UV-C was reported by the World Health Organization to be the most damaging spectrum of sunlight to full thickness skin (9,22). Thus, UV-C at the intensities used by traditional germicidal equipment is only safe for pre- and post-operatively use when humans are not at risk for exposure. Skin, however, is differentially affected by the various wavelengths of UV light. UV light can be subdivided into the UV-C spectrum (100-280 nm), UV-B spectrum (280-315 nm), and UV-A spectrum (315-400 nm) (9). In contrast to UV-C, longer wavelengths of the UV spectrum, particularly UV-A, are able to penetrate deeper layers of the skin (e.g., dermis) (9,23).

Similar to UV-C, the UV-A spectrum also has antimicrobial properties (24,25). UV-A has been shown to indirectly kill cells through the accumulation of free radicals (24,26,27). UV-A is abundantly present in sunlight and can be germicidal at doses that do not pose a risk to humans (25,26). Thus, UV-A could conceivably be a safe option for disinfection in healthcare settings. Currently, a UV-A device for use in the operating room is under development and is already shown to successfully reduce pathogens on medical equipment (25). Industry standards for photobiological safety of germicidal lighting limit human exposure to UV-A to an irradiance of 10 W/m^2^ (28–30). Previous studies using UV-A devices similar to those used in this study found that when irradiance at the head or eye was limited to 10 W/m^2^, the irradiance at heights roughly equivalent to those of operating tables were approximately 3 W/m^2^ (25,31). Accordingly, we hypothesized that appropriately dosed levels of continuous UV-A irradiance could be used intraoperatively with an acceptable safety profile for both patients and medical personnel. The safety of using UV-A light in the operating room is not known. Thus, our study was required to begin defining the safety profile of intraoperative UV-A exposure.

## METHODS

The UV light devices were provided by GE Current, a Daintree company (East Cleveland, OH). UV-C and UV-A intensities (W/m^2^) were measured using a UV light meter (Digi-Sense). All animal experiments were conducted under institutionally approved protocols (IACUC-07733 for *Sus domesticus* and IACUC-21885 for *Mus musculus*).

### Immunoblotting for DNA damange and DNA repair proteins

HEK293 (human embryonic kidney cells) were plated at a density of 10^5^ cells and incubated overnight at 37°C and 5% CO^2^ to reach 80% confluence in T-75 flasks. They were subsequently exposed to either 1 or 2 hours of either UV-A light at 30 W/m^2^ (10 times what the UV-A irradiance may be in a typical application) or UV-C light at 0.3 W/m^2^. Additional HEK293 cells were plated in the same fashion and were then exposed to either room LED (light-emitting diode) light as a negative control or 10 Gy of gamma radiation over 10 minutes as a positive control. Whole cell lysates were collected 30 minutes after light exposure. Proteins known to be activated via phosphorylation after DNA damage or involved in DNA repair were compared between conditions (pH2A.X and pCHK1). Antibody conditions to detect each protein were: H2A.X (1:500, Proteintech 10856-1-AP), pH2A.X (1:500, Cell Signaling Technology #9718), pCHK1 (1:500, Cell Signaling Technology #2348), and the loading control GAPDH (1:1000, Santa Cruz sc-47724).

### Alkaline Comet Assay to detect DNA damage

HEK293 (human embryonic kidney cells) and WI-38 (human lung fibroblasts) were plated to 10^5^ cells and incubated overnight at 37°C and 5% CO^2^ to reach 80% confluence in T-75 flasks. They were subsequently exposed for 2 hours to either room light, UV-A light at 30 W/m^2^, or UV-C light at 0.3 W/m^2^. A CometAssay kit (Trevigen 4250-050-K) was used to process cells. SYBR Green (ThermoFisher S7563) was used for nuclear staining. Comet Assay IV (Instem) software was used to measure tail length and momentum and to calculate % tail DNA. A total of 300 cells were measured for each condition. The Kruskal-Wallis test was used to analyze comet tail length and % tail DNA using IBM SPSS Statistics for Windows, version 25 (IBM Corp, Armonk, NY). Statistical significance was defined as P < 0.05.

### Immunohistochemistry for DNA damage-related photoproducts

4-week-old CD-1 mice (equal numbers of males and females) were anesthetized with isoflurane, dehaired (Nair Hair Removal Lotion), and then exposed to either room light, UV-A light at 30 W/m^2^, or UV-C light at 0.3 W/m^2^ for a total of 2 hours. Mice were immediately sacrificed after exposure. The tissues harvested for staining included the liver, and small bowel. Tissues were fixed in 10% formalin, paraffin-embedded, and then sectioned (UAB Animal Pathology Lab). Retrieved slides were deparaffinized by hydrating tissues in Xylene (5 min x2), 100% ethanol (5 min x2), 95% ethanol (5 min x1), 75% ethanol (5 min x1), and then distilled water (2 minutes) followed by placement in pre-warmed citrate antigen retrieval buffer (BioGenex #HK086-9K) and pressurized in a pressure cooker for 10 minutes. The slides were subsequently cooled for 15 minutes prior to further processing in 3% hydrogen peroxide in the dark (10 minutes). Slides were gently washed in phosphate buffered saline (PBS) prior to blocking with the blocking reagent from the mouse-on-mouse polymer IHC Kit (Abcam ab269452). The tissues were then incubated overnight with the primary antibody at 4°C. Markers evaluated includes cyclobutene pyrimidine dimers (1:1000, CPD, CosmoBio CAC-NM-DND-001) and pyrimidine-pyrimidone (6-4) photoproducts (1:1000, 6-4 PP, CosmoBio CAC-NM-DND-002). Afterwards, the slides were PBS washed prior to incubating with mouse secondary antibody (1:500, Invitrogen A16070) for 1 hour. The slides were then washed again before incubating with HRP polymer detector reagent from the same mouse-on-mouse IHC kit for 20 min. The slides were then stained for 2 minutes 30 seconds with a DAB substrate kit (Abcam ab 64238) and washed in tap water twice prior to incubating in tap water for 2 hours prior to cover slipping.

### Porcine model for projection of UV-A reflection

A porcine model was used to measure the reflected irradiance of UV-A light. The animal was anesthetized per standardized protocol in a supine position. An overhanging UV-A light fixture was set to provide irradiance of 30 W/m^2^ at the level of the skin (Figure 5A-B). Irradiance measurements were obtained at 3 positions 1 foot away from the animal at 3 varying heights (Figure 5C-D). UV-A irradiances were measured in triplicates at the designated locations and final values were averaged for two conditions: closed skin and open abdomen.

### Personal Protective Equipment (PPE) model for UV-A protection

To evaluate the level of UV-A protection provided to surgical staff by typical equipment, small samples approximately 50-75mm square were cut from samples of surgical Personal Protective Equipment (PPE) used in the operating environment. Between 4 and 6 samples were taken for each type of PPE. In instances where the PPE was composed of multiple materials, samples were taken of each material type and evaluated separately. A UV/visible spectrophotometer (Lambda 950, PerkinElmer, Inc.) was used to measure spectral transmission of these samples over the full UV-A wavelength range (315-400nm). To evaluate the transmission from the UV-A disinfection device used in this study, an average of spectral transmission over the range of 360-370nm was taken.

## RESULTS

### UV-A light induces no more DNA damage than standard room lighting at the cellular level

To investigate potential effects of UV-A versus UV-C light on DNA damage and DNA reapair mechanisms, we looked at changes in intracellular markers corresponding to damage and repair pathways. Specifically, we assessed protein-level expression of phospho-pH2A.X and phospho-CHK1 (pCHK1) (32,33) (Figure 1). Cells exposed to 10 minutes of 10 Gy radiation served as a positive control. After 1 hour of exposing cells to UV-A light, there was less phosphorylation of histone H2A.X (pH2A.X) compared to normal LED light (control) and UV-C light. After 2 hours of exposure, only cells that received UV-C light showed an increase in phosphorylation of CHK1 (pCHK1). The activation of CHK1, detected by the phosphorylation of CHK1 (pCHK1), was only induced in cells exposed to UV-C for 2 hours and the positive control, cells exposed to 10 Gy gamma irradiation. This data suggests that at the cellular level UV-A exposure does not significantly induce DNA damage repair pathways.

**Figure 1:**
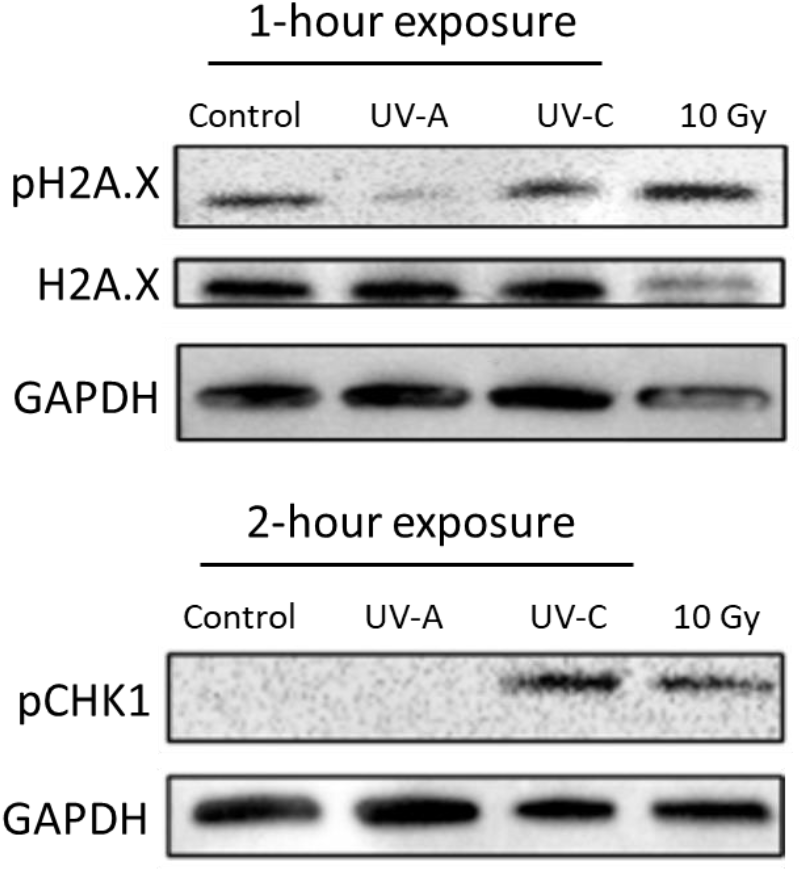
Western blot comparing markers of DNA repair. After 1 hour of exposing cells to UV-A light, there was less phosphorylation of histone H2A.X (pH2A.X) compared to normal LED light (control) and UV-C light. After 2 hours of exposure, only cells that received UV-C light showed an increase in phosphorylation of CHK1 (pCHK1). Cells exposed to 10 Gy of gamma radiation over 10 minutes served as a positive control.

Further investigation into the impact of UV-A versus UV-C light on DNA damage was done by measuring the amount of double stranded DNA breaks using a Comet Assay. Double stranded DNA breaks were quantified by (1) comet tail length and (2) %tail DNA. In two different cell lines (HEK293 and WI-38), UV-C exposure caused significantly greater tail lengths and %tail DNA compared to a dose of UV-A that was 10-fold higher (Figure 2). This result indicates a higher level of DNA damage, in terms of double stranded DNA breaks, in cells exposed to UV-C as compared to UV-A light. In both cell lines, the normal room light condition demonstrated greater %tail DNA than UV-A exposure, suggesting that UV-A light does not induce any more double-stranded DNA breaks than normal room light condition alone.

**Figure 2:**
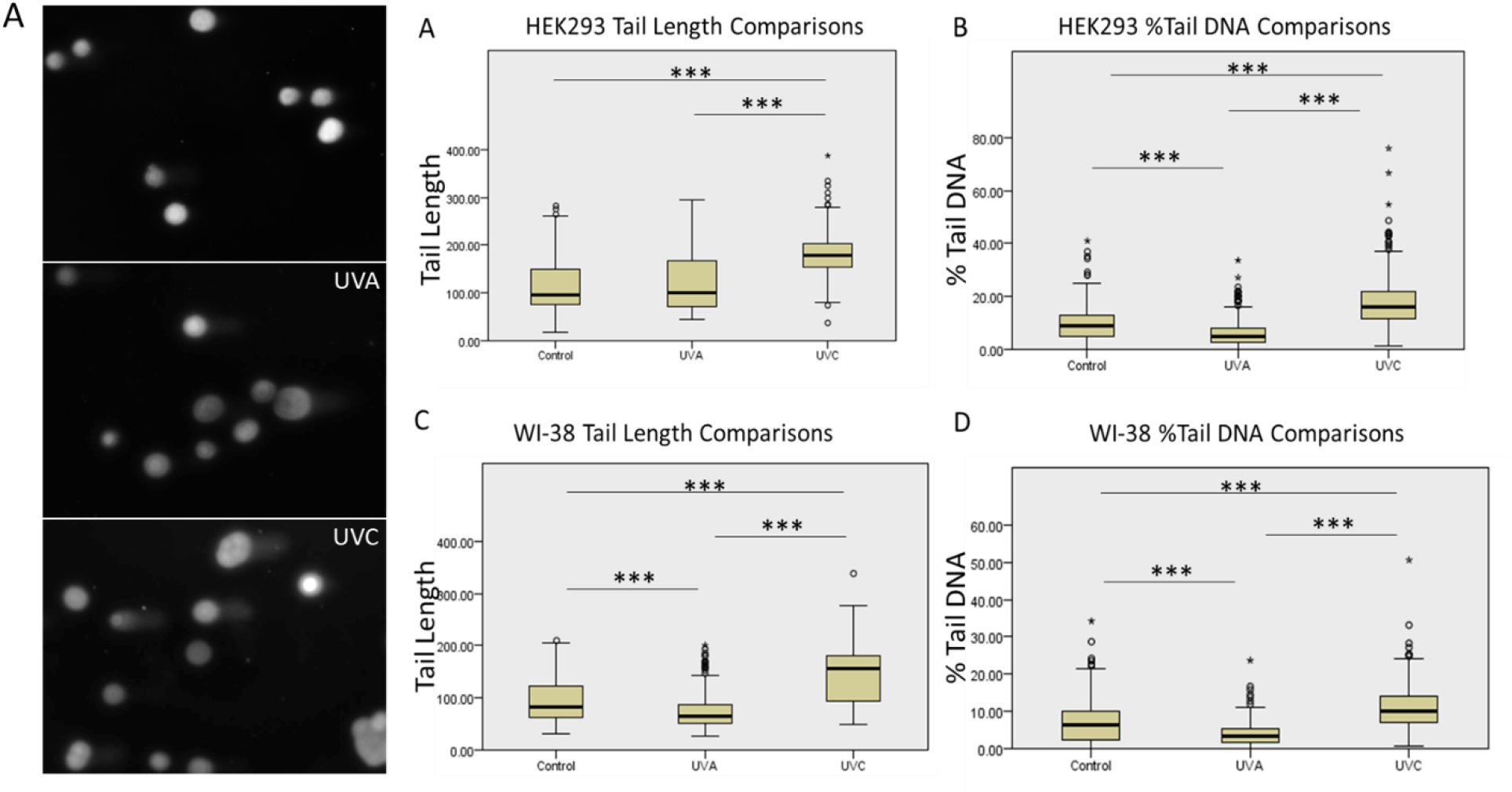
DNA damage after UV-A/UV-C light exposure assessed by a comet assay. **(A)** Representative images of comet assay DNA damage in HEK293 cells exposed to either room light, UV-A, or UV-C light. Comparisons in tail length in **(B)** HEK293 and **(D)** WI-38 cells. Comparisons in %tail DNA in **(C)** HEK293 and **(E)** WI-38. N = 300, *** indicates a *p* < 0.001.

### Intraoperative UV-A exposure does not produce DNA damage-related photoproducts in tissue

DNA damage caused by UV light exposure creates cyclobutyl pyrimidine dimers (CPD) and pyrimidine (6-4) pyrimidone photoproducts (6-4PP) (9,16,17). These photoproducts accumulate in the nucleus and can lead to cell death. We sought to determine if tissues intraoperatively exposed to UV-A light produced CPD and 6-4PP, in comparison to either intraoperative UV-C or the normal room light condition. We used a murine model to administer each light group to the serosal surface of the abdominal viscera. Immunohistochemistry was used to detect the presence of CPD and 6-4PP in nuclei after light exposure in the small intestine (Figure 3) and liver (Figure 4). Small intestinal and liver tissue exposed to either room light or UV-A light did not show any positive nuclear staining for either CPD or 6-4PP.

**Figure 3:**
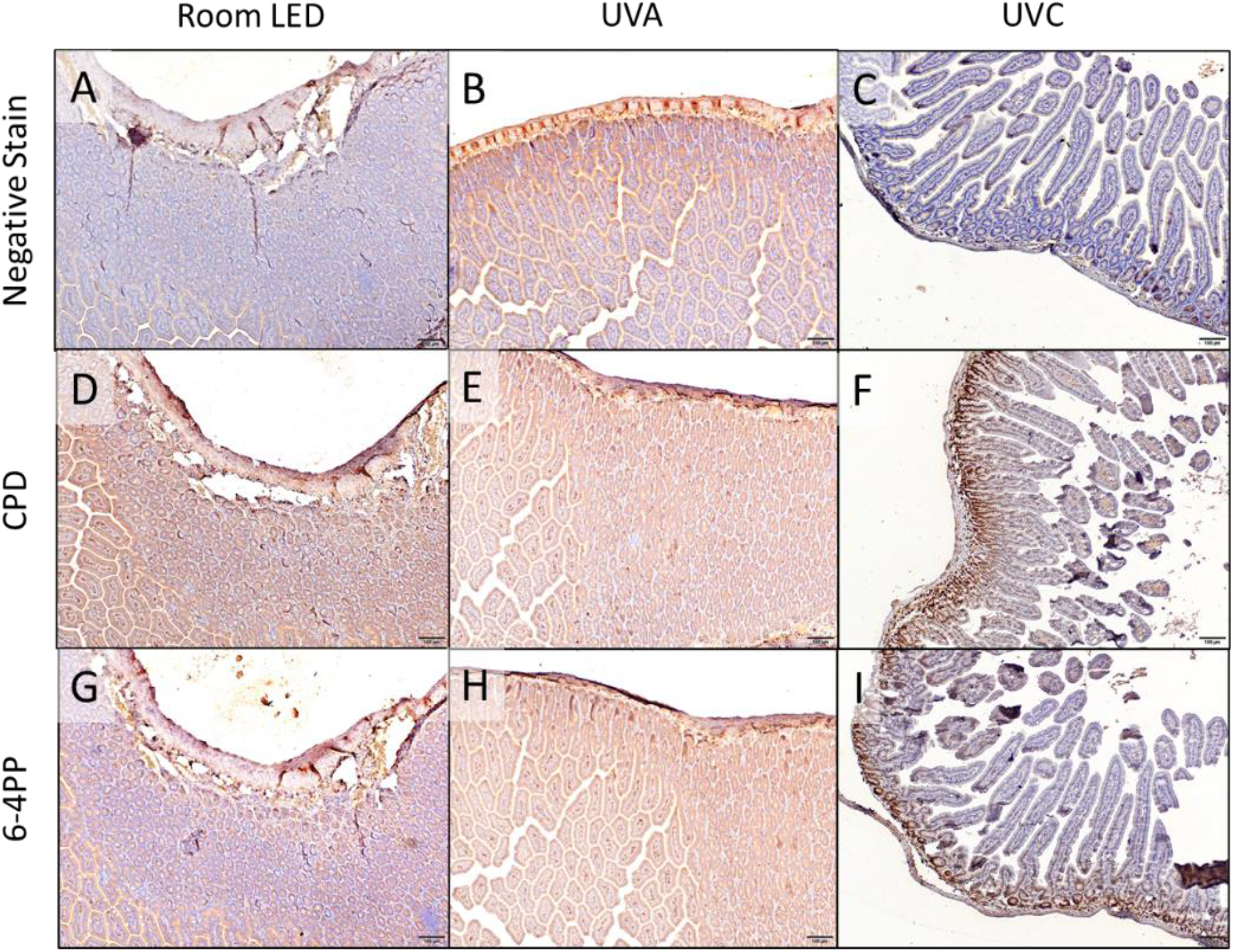
Comparison of DNA damage photoproducts CPD and 6-4PP in the small intestines after either room light, UV-A, or UV-C light exposure. Negative stain controls for small intestinal tissue exposed to **(A)** room light, **(B)** UV-A, or **(C)** UV-C light. Positive nuclear staining for CPD and 6-4PP was confirmed in small intestinal tissue exposed to UV-C light **(F, I)**. Small intestinal tissue exposed to either room light or UV-A light did not show any positive nuclear staining for either CPD **(D, E)** or 6-4PP **(G-H)**. Scale bar = 50 µM.

**Figure 4:**
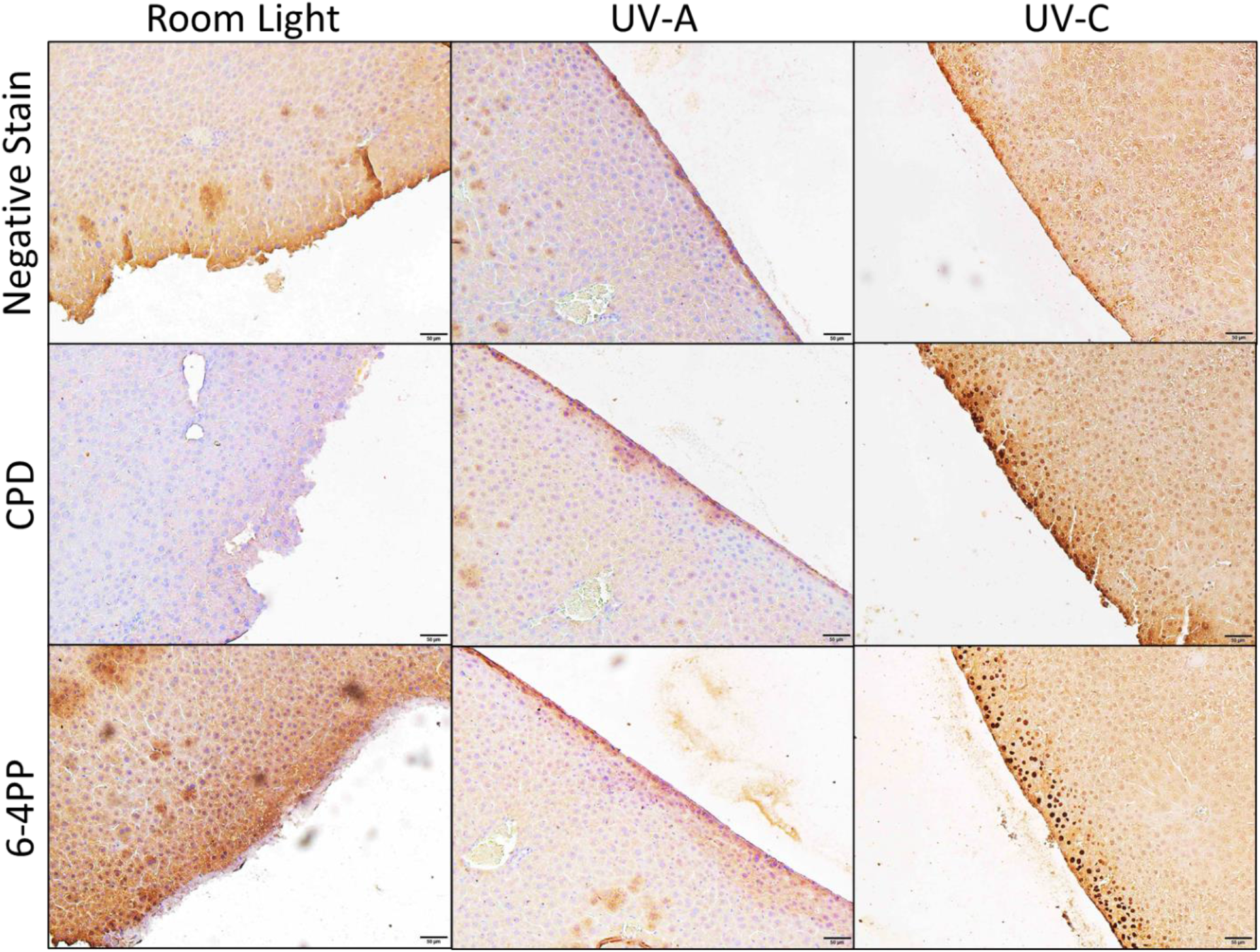
Comparison of DNA damage photoproducts CPD and 6-4PP in the liver after either room light, UV-A, or UV-C light exposure. Negative stain controls for liver tissue exposed to room light, UV-A, or UV-C light. Positive nuclear staining for CPD and 6-4PP was confirmed in liver tissue exposed to UV-C light. Liver tissue exposed to either room light or UV-A light did not show any positive nuclear staining for either CPD or 6-4PP. Scale bar = 50 µM.

### Intraoperative UV-A produces a safe level of reflection on surgical personnel

to determine the reflective irradiance exposure of surgical personnel by UV-A, we employed a porcine model of abdominal surgery (Figure 5). We showed that the greatest amount of reflected UV-A irradiance was 20-fold less than the patient’s exposure (0.15 mW/cm^2^ versus 3.0 mW/cm^2^). The pattern of reflected UV-A light irradiance did not differ between the surgeon and first assist positions, where the surgeon stood 12 inches to the left of the pig and the first assist stood 12 inches to the right (Figure 5C). We then measured the amount of UV-A reflection at different personnel heights (60, 68, 77 inches tall) for the surgeon, first assist, and a surgical technican position (Figure 5D). We found that the visceral reflection of UV-A light had a higher irradiance at shorter personnel heights compared to skin exposure, but the visceral reflected irradiance remained lower at taller personnel heights (Figure 5E).

**Figure 5:**
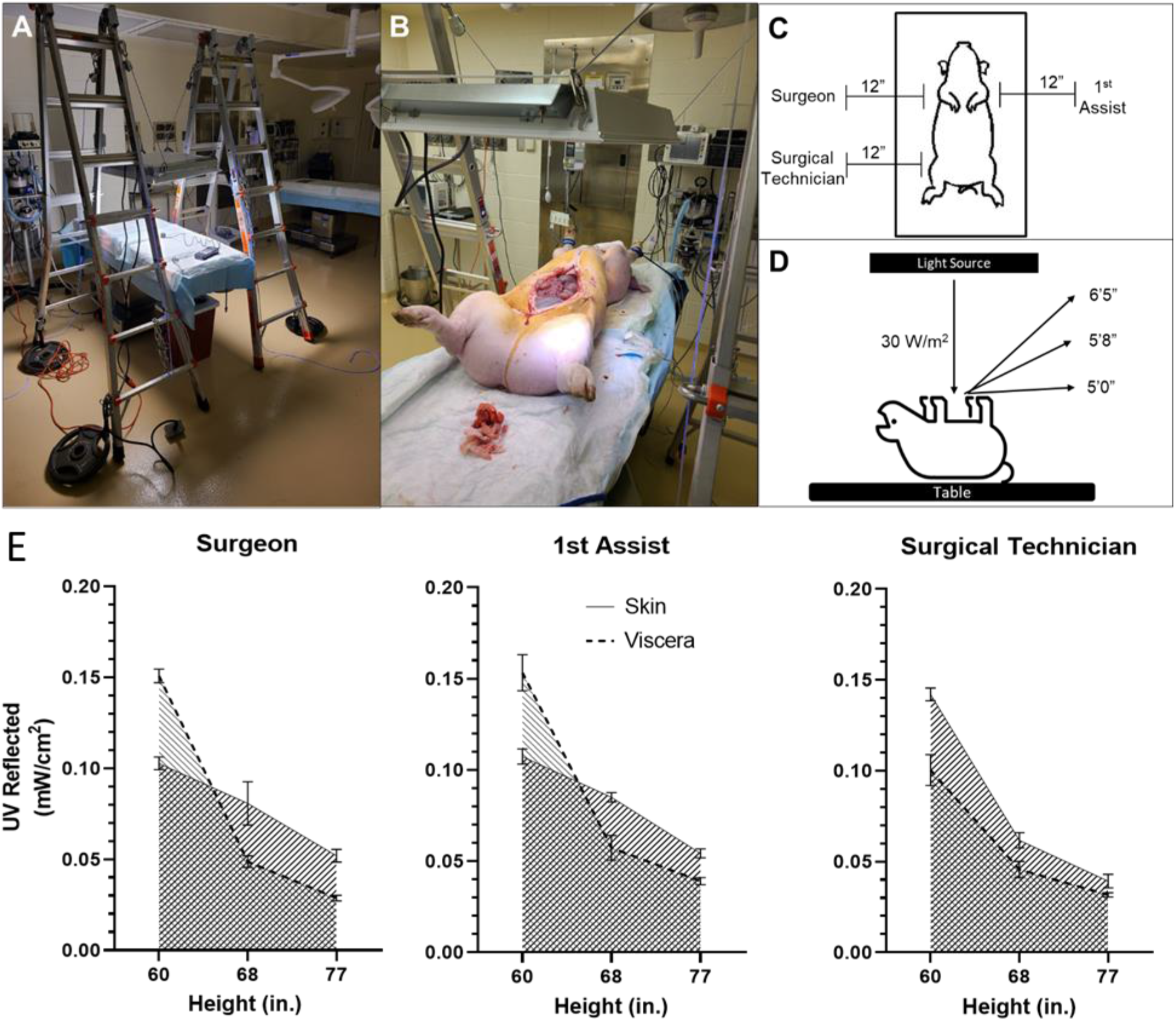
Model of provider exposure to UV light. **(A)** Overhanging light to achieve UV-A irradiance of 30 W/m^2^. **(B)** Light source overlying pig with adjustable light source using pulley/lever system. **(C)** Operative team positions around the porcine subject for reflected UV-A measurements. **(D)** Height variables for light measurements. **(E)** Comparison of reflected UV-A irradiance measured at the surgeon, 1^st^ assist, surgical technician locations. Solid line marks reflected irradiance based off skin only measurements. Dotted line represents reflected irradiance based off visceral intensities.

### Analysis of UV-A transmission, transmittance, and absorbance on surgical PPE

To evaluate an additional safety consideration of using UV-A during surgery, we measured the transmission of UV-A light through surgical PPE. Transmittance (transmitted ratio) and absorbance (-log_10_ of transmittance) were calculated from the measured transmission. We evaluated twelve different types of common surgical PPE, including eyeglasses, surgeon’s caps, surgical masks, and surgical gloves. Brand information and reference IDs for each type of PPE are provided in Supplemental Table 1 (Table S1). Percent transmission, transmittance (τ, where τ=%T/100), and absorbance (A, where A=-log_10_τ) for the range of 360-370 nm are provided in Table 1 for the measured PPE. All surgical PPE measured in this study absorbed UV-A at varying levels. Surgical gloves provided the highest level of UV-A attenuation among the measured items, transmitting less than 10% of UV-A in the 360-370nm range. All opaque PPE transmitted 24% or less of these wavelengths, while transparent or mesh fabric PPE transmitted up to 90%.

**Table 1:** Transmission, transmittance, and absorbance of surgical PPE in the range of 360-370nm

| **Measured PPE** | **Percent Transmission (%T) of UV-A (360-370nm, %)** | **Transmittance ($\tau$) of UV-A (360-370nm)** | **Absorbance (A) of UV-A (360-370nm)** |
| --- | --- | --- | --- |
| Drape leggings (clear plastic part) | 90 | 0.90 | 0.04 |
| Drape leggings (fabric part) | 10 | 0.10 | 1.00 |
| Eyeglasses | 80 | 0.80 | 0.10 |
| Surgeons Cap (white part) | 73 | 0.73 | 0.13 |
| Surgeons Cap (Blue part) | 19 | 0.19 | 0.72 |
| Bouffant Cap | 64 | 0.64 | 0.19 |
| Light Blue Gown | 24 | 0.24 | 0.62 |
| Dark Blue Gown | 15 | 0.15 | 0.83 |
| Surgical Mask (blue) | 18 | 0.18 | 0.74 |
| Surgical Mask (green) | 17 | 0.17 | 0.78 |
| Drape | 11 | 0.11 | 0.94 |
| Polyisoprene Surgical Glove | 9 | 0.09 | 1.02 |

## DISCUSSION

Surgical site infections (SSIs) pose an ongoing threat to patients’ physical and mental health, and the costs of complications associated with SSIs continue to burden the healthcare system. UV light has historically been used as a disinfection agent for inanimate objects. Theoretically, UV light could be used a technique to reduce SSIs by maintaining a sterile surgical field, although the safety of intraoperative UV-A light exposure must first be evaluated. For example, UV-C light is used in healthcare settings to disinfect surfaces in the operating room, hospital bed rooms, and ambulances (9–12). However, UV-C over-exposure is associated with health risks (9,21,22). A safer alternative to typical UV-C light disinfection products could be devices that use low levels of UV-A light. Previous studies have shown that UV-A light is germicidal and is capable of reducing pathogens on steel surfaces (25). UV-A exposure from an overhead light source significantly reduced the amount of pathogenic microorganisms on medical equipment (25).

This study aimed to evaluate the safety of using intraoperative UV-A light. Previously, the safety profile of using UV-A light during an operation was unknown, as all commonly used photobiological standards are based on studies of exposed skin or eyes and not internal organs. We found exposure to UV-A light did not cause significant DNA damage. We tested for DNA damage in cells exposed to UV-A light by using a comet assay to detect DNA double stranded breaks, along with western blot to detect expression of DNA damage-related proteins (32–34). In further investigation of potential UV-A-induced DNA damage, we did not find any photoproducts (CPD or 6-4PP) formed in the nuclei of internal organ tissue that was exposed to UV-A. These photoproducts are specifically produced by UV exposure, making them a reliable readout for UV-induced DNA damage (35). Taken together, these data suggest that UV-A light does not induce a significant amount DNA damage in cells or tissues.

Our results also showed UV-A reflection in operating room remained at low levels, thus not posing a health risk to surgical personnel. We found that the visceral reflection of UV-A light had a higher irradiance at shorter personnel heights compared to skin exposure, but the visceral reflected irradiance remained lower at taller personnel heights. At the position of the surgical technician, there was greater reflective irradiance with the skin exposure at all heights compared to visceral exposure. These differences may be due to differences in UV-A reflectance, specularity, or overall shape between skin and viscera. Furthermore, the amount of reflected UV-A surgical personnel experience when UV-A was adminstered overheard during a procedure was less than 20-fold of what was experienced by the patient. The industry photobiological standards limit continuous UV-A exposure to 10 W/m^2^. If a system is designed to limit exposure at the patient to this level, the surgical staff will experience over an order of magnitude less irradiance than the patient. Furthermore, it is also important to consider reflected UV-A onto surgical staff because the eye is more sensitive to UV-A than the skin. We would expect, if using an overhead UV-A disinfection light, that the highest irradiance would be at the top of the surgeon’s head and at a much lower irradiance at the eye (looking downward at the table or horizontally). Overall, the amount of UV-A irradiance reflected onto surgical personnel was below industry photobiological standards limits of 10 W/m^2^ for continuous UV-A exposure, which indicated that overhead UV-A exposure during an operation did not pose a health risk.

In our study of UV-A absorption by surgical PPE, each item measured absorbed some UV-A. In practice, a ceiling-mounted UV-A disinfection system that abides by accepted industry photobiological safety standards would not expose humans to levels of UV-A that may be hazardous (28,29). While UV-A-absorbing PPE is not necessary, using such may increase confidence in the safety of a UV-A by surgical staff who would be frequently exposed to germicidal wavelengths.

Although we examined multiple aspects of UV-related DNA damage and intraoperative UV-A reflection, the studies described herein had several limitations. First, we did not investigate all molecular pathways of DNA damage. Implementation of overhead UV-A devices in procedures would naturally require that the other equipment be adequately integrated. In future studies, it will be imperative to investigate the potential ability of UV-A light to reduce SSIs, as confirmation and successful implementation of this modality could greatly benefit patient care. Overall, our findings suggest that UV-A light can safely be used in the operating room.

## Author Contributions

### Author contributions

R.Z.P performed the comet assay and the murine study looking at DNA damage-related photoproducts via immunohistochemistry. B.H. performed the porcine preclinical safety study. R.G. and M.M. wrote the manuscript. K.J.B. provided the results of the PPE safety study and edited the sections of the manuscript related to description of the devices and safety. All authors revised the manuscript.

